# Progress in quickly finding orthologs as reciprocal best hits

**DOI:** 10.1101/2020.05.04.077222

**Authors:** Julie E Hernández-Salmerón, Gabriel Moreno-Hagelsieb

## Abstract

**Introduction:** Finding orthologs remains an important bottleneck in comparative genomics analyses. While the authors of software for the quick comparison of protein sequences evaluate the speed of their software and compare their results against the most usual software for the task, it is not common for them to evaluate their software for more particular uses, such as finding orthologs as reciprocal best hits (RBH). Here we compared RBH results, between prokaryotic genomes, obtained using software that runs faster than blastp. Namely, lastal, diamond, and MMseqs2.

**Results:** We found that lastal required the least time to produce results. However, it yielded fewer results than any other program when comparing evolutionarily distant genomes. The program producing the most similar number of RBH as blastp was MMseqs2. This program also resulted in the lowest error estimates among the programs tested. The results with diamond were very close to those obtained with MMseqs2, with diamond running faster. Our results suggest that the best of the programs tested was diamond, ran with the “sensitive” option, which took 7% of the time as blastp to run, and produced results with lower error rates than blastp.

**Availability:** A program to obtain reciprocal best hits using the software we tested is maintained at https://github.com/Computational-conSequences/SequenceTools

## 1 Introduction

Finding orthologs is a central step in comparative genomics and represents a central concept in evolution. Orthologs are defined as characters that diverge after a speciation event [1]. In other words, if the characters are genes, then they are thought of as the same genes in different species. It is expected that they typically conserve their original function, an inference that has been supported by several lines of evidence [2, 3, 4, 5].

Efforts in standardizing orthology inference methods remain in constant evaluation, with over forty web services available to the community [6, 7]. Few of these methods are based on phylogenetic analyses (tree-based approach), which, despite expected to be the most accurate, tend to be computationally intensive and impractical for big databases [8, 9]. Some methods employ pairwise sequence similarity comparisons (graph-based methods) that have been successful to implement, such as the clusters of orthologous groups (COGs) database [10, 11]. However, researchers require having their own sets of orthologs, since genome sequencing has become a commonly available technology.

Perhaps the most common approach, or working definition, of orthology, is Reciprocal Best Hits (RBH), which is a simple method that has shown low false-positive rates and ease of implementation [12, 9, 13]. Essentially, if gene *a* in genome *A* finds gene *b* as its best, highest-scoring, match in genome *B*; and gene *b* finds gene *a* as its best match in genome *A*, they are RBH and thus inferred to be orthologs. The most commonly used program for finding sequence matches for finding RBH is blastp [14]. This program was chosen for being the fastest program available at the time when comparative genomics started. However, the amount of sequences to analyze continues to grow exponentially, making the speed of blastp comparisons too slow for the increasing demand of sequence analysis.

When authors introduce a software suite for sequence comparison, they often compare the speed and overall performance of their software compared to blastp. Since speed tends to come at a cost in sensitivity and accuracy, the author reports might include differences in performance in overall sequence comparison and number of detected sequences. However, more specialized usages, like finding orthologs as RBH might be affected differently compared to overall performance. Thus, it becomes necessary to test the adequacy of the software in particular tasks. Accordingly, prior work compared the performance of three fast programs against blastp [15]. The programs tested were blat [16], ublast [17] and lastal [18]. Since then, two somewhat recently developed programs for sequence comparison include diamond [19] and MMseqs2 [20] (from now on mmseqs). Here we use lastal as a reference to the previous report [15], where lastal was the program producing the most-similar-to-blastp results, and test the performance of these two new programs, diamond and mmseqs, for obtaining RBH.

## 2 Methods

The annotated protein sequences of 3,312 non-redundant genomes were used as subjects for comparisons against the genome of *Escherichia coli* K-12 MG1655 (GCF_000005845) as the query. The non-redundant representatives were selected from 1̃6,000 complete genomes available at NCBI’s refSeq genome database [21] by January 2020. To select these genomes, we clustered them using a trinucleotide DNA signature [22], with a δ-distance cutoff of 0.04 as described previously [23], resulting in 3312 clusters. A distance that roughly corresponds to a “species” level. We took one genome per cluster.

Four programs were used to perform protein sequence comparisons: (1) blastp version 2.10.0+ [14] (ftp://ftp.ncbi.nlm.nih.gov/blast/executables/blast+/), (2) lastal version 1045 [18] (http://last.cbrc.jp), (3) diamond version 0.9.30 [19] (https://github.com/bbuchfink/diamond), and (4) mmseqs version 11-e1a1c [20] (https://github.com/soedinglab/MMseqs2).

To compare times, the query *versus* targets comparisons were run one by one, in the same computer, with no other programs running at the same time. Times were obtained by using the unix “time” command. This command reports user, cpu and real times. The plotted times were the real times.

To find Reciprocal Best Hits (RBH), we wrote a program, getRBH.pl, to standardize the options and outputs from the different sequence comparison programs, following the options previously described by Ward and Moreno-Hagelsieb [15], the e-value threshold was 1 × 10^−6^ (1e-6), coverage of 60% of the shortest protein in the alignment, as well as soft-masking and Smith-Waterman alignment, when available (diamond does not have soft masking, thus it was run with no masking). The latter two previously found to improve the finding of RBH [12]. Since all four programs can work with their own target (also called subject) databases, we built databases for all programs.

Genomic Similarity Scores (*GSS*) were calculated from blastp results as the sum of the bit scores of all reciprocal best hits *(compScore)* divided by the bit scores of each corresponding ortholog against itself *(selfScore)*, as described in [23] as *GSSa*.

Error estimates in detecting orthologs were calculated following conservation of gene order as means for testing quality. For this, we analyzed adjacent homologs genes, assuming that a pair of conserved genes in the target genome will be both orthologs to the corresponding pair of genes in the query genome [12].

## 3 Results and Discussion

### 3.1 Runtimes

The computing speeds for finding homologs were timed for each program relative to blastp. Of all the programs tested, lastal was the fastest, finding the corresponding homologous proteins in 1.5% of the time spent by blastp (Fig. 1).

**Figure 1.**
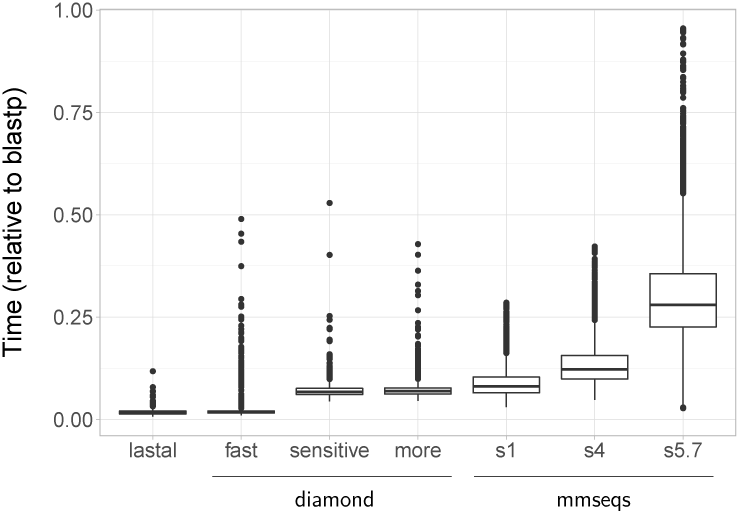
Difference in speed obtaining pairwise alignments between the programs tested. The times plotted are the “real” times, as measured by the *time* UNIX command, relative to blastp. The fastest of the three programs we tested was lastal. Both diamond and mmseqs were tested with different sensitivity options.

Both diamond and mmseqs can be run with different sensitivity options. The available sensitivity modes for diamond are “fast”, “sensitive” and “more-sensitive”. We did not find much difference in the runtimes for the last two modes, which took around 7% of the time it took blastp to run. The “fast” mode took slightly less than 2% of the blastp time to run, thus being the closest one to the lastal time. The sensitivity options for mmseqs tested were 1, 4, and 5.7. 1 and 4 were chosen because both were used in the article presenting the software [20], while 5.7 is the default option.

Note that we ran mmseqs with the “easy-search” function. This function produces any output format desired without much user intervention. The easy-search function accepts the target either in plain fasta format, or as a formatted database, but the query has to remain in plain fasta format. Another way to produce pairwise alignments with mmseqs would use the “search” function instead of easy-search. The search function requires databases built for both, query and target. The results of the search function is also in database format. This database can then be used to extract results into other output formats as necessary.

The mmseqs software can also precompute indexes for its databases. We decided not to built indexes because they take very long to be built and use too much space. For example, the database for the longest of the subject genomes (16,040,666 bp) used 5.7M of space, which increased to 898M when building the index. Runtimes might vary if the user preferred to build a database index and use the search function instead of the easy-search one.

Note that mmseqs has a “rbh” function, with a future version offering an “easy-rbh” function which should take care of producing a table without much user intervention (Martin Steinegger, personnal communication). However, we decided not to use the rbh function because we preferred to keep control of the parameters used to produce RBH.

### 3.2 Reciprocal best hits

The proportion of reciprocal best hits found using the faster programs was evaluated with respect to blastp (Fig. 2, 3). Our results showed a very similar proportion of RBH as those obtained with blastp when genomes are more similar to *E. coli* K12 MG1655. As the similarity decreased, so did the proportion of RBH found as compared against those found by blastp.

**Figure 2.**
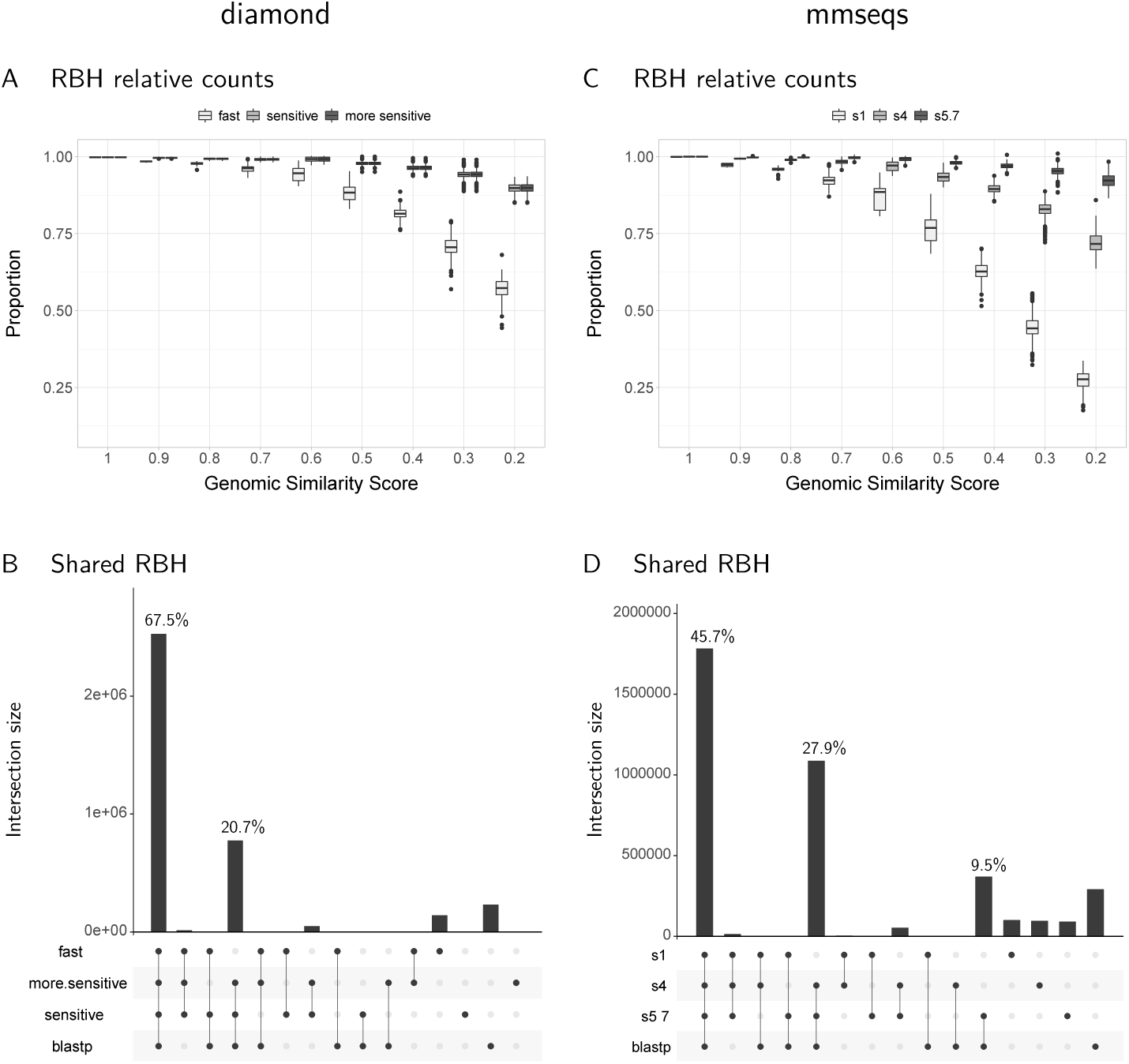
Reciprocal best hits found by diamond and mmseqs. A,C: The proportion of RBH found is comparable to those found by blastp when the genomes involved are very similar (high Genomic Similarity Scores, GSS). This proportion reduces with the GSS. The effect of sensitivity options, as expected, is a reduced proportion of RBH found when setting for low sensitivity. For diamond, the top two sensitivity modes produced very similar results. B,D: Most of the RBH found correspond to RBH found by blastp.

**Figure 3.**
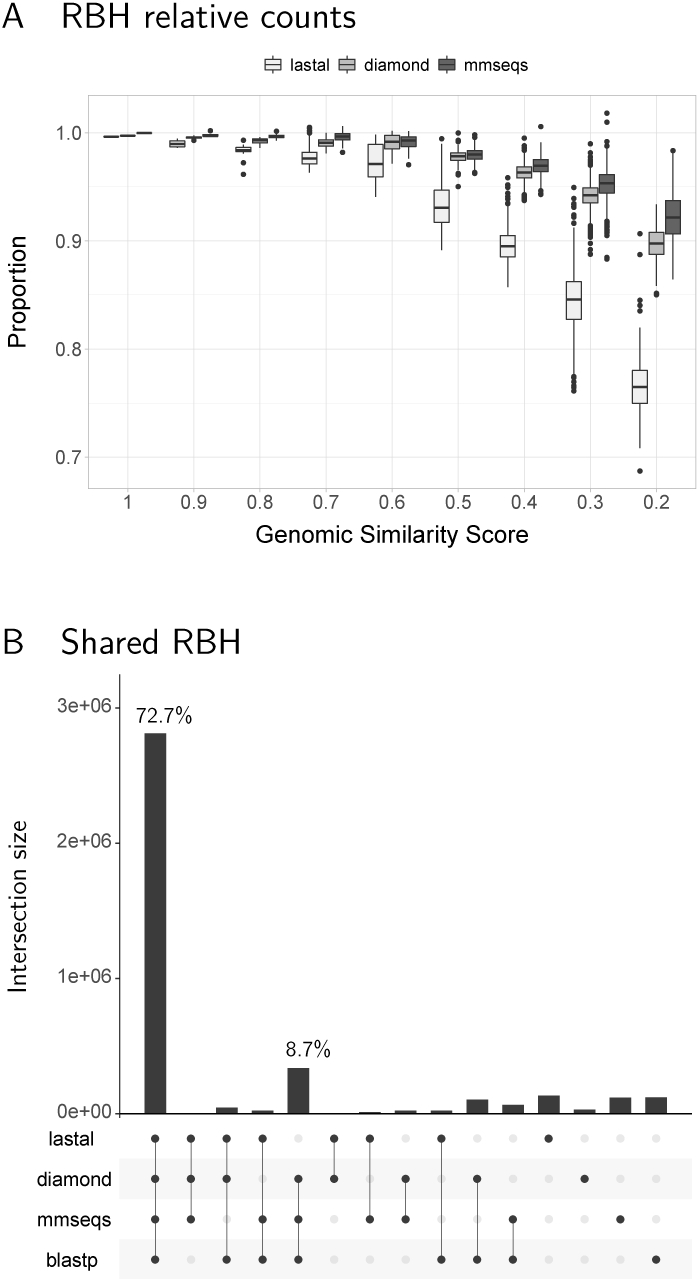
Reciprocal best hits found by all programs. A: The proportion of RBH obtained with the fast programs, relative to the number obtained with blastp, is very similar when genomes are very similar to the query genome (here *E. coli* K12 MG1655). As genomes become less similar, the proportion falls. Both diamond and mmseqs showed improved proportions at low overall similarity compared to lastal. B: Close to 73% of the RBH detected are shared by all programs. Both diamond and mmseqs share the most RBH with blastp.

The different sensitivity options resulted in different proportions of RBH found by either diamond or mmseqs (Fig. 2). As would be expected, the least sensitive options resulted in fewer RBH compared to blastp. In the case of diamond, the proportions found using the fast setting dropped noticeably below those detected with the other options (Fig. 2A). Furthermore, the sensitive and more sensitive settings had the highest RBH in common with blastp for a total of 88.4% (67.7% + 20.7%) (Fig. 2B). Thus, the fast option did not represent the best option for detecting RBH with diamond. It also appears that the more-sensitive mode has no advantage over the sensitive option. Since the sensitive option is the main one discussed in the diamond article [19], we decided to use this option for further analyses.

In the case of mmseqs, the sensitivity options tested produced very different results (Fig. 2C). Again, the top options, 4 and 5.7, shared the most results with blastp (Fig. 2D), though only amounting to a combined 73.6% (45.7% + 27.9%). The 5.7 option produced the best results, with 9.5% more RBH shared with blastp. We thus chose this option for further analyses.

We thus compared the results obtained at the above chosen sensitivity options with those obtained with lastal. Both diamond and mmseqs detected a higher proportion of RBH than lastal (Fig. 3A). This was true even at the lowest GSS values, meaning that even in the worst cases, mmseqs and diamond would not miss more than 10% of the RBH that would be produced by blastp. The mmseqs results were the best in this sense. However, this option ran in around 28% of the time as blastp, while the diamond option chosen takes around 7% of the time used by blastp.

Around 73% of all RBH were detected by all programs (Fig. 3B). The second most important intersection shows that diamond and mmseqs shared the majority of RBH with blastp (making up a total of 81.4%). Of the total RBH, 3.2% were only found by blastp, while 7.4% was detected by the fast programs but not by blastp. Unlike our previous analysis [15], which showed higher percentages of RBH only detected by blastp, similar numbers of RBH were found by both mmseqs and lastal (3.1% and 3.5% respectively). These percentages seem to represent differences in sensitivity, with these cases being either differentially detected true positives, or false positives.

### 3.3 Error estimates

Despite genomic rearrangements and horizontal gene transfer result in divergence of gene order, remnants of highly conserved regions are preserved even between the genomes of evolutionarily distant organisms [24, 25, 26]. Thus, despite conservation of adjacency is a very limited source for correction of misidentified orthologs, the few pairs of adjacently conserved genes can be used to estimate error rates by extrapolation [12]. The estimate consists on finding pairs of adjacently conserved gene pairs. If both genes in a target pair are orthologs to the corresponding gene pair in the query genome, they are inferred to be correct. Errors would consist on conserved pairs where only one of the genes is detected as an ortholog, with the remaining gene being evidence of an ortholog mistaken for a paralog. The formula for error rates is: *E* = *p/p* + *o*, where *p* stands for number of paralogs in conserved pairs, and *o* for the number of orthologs in conserved pairs [12].

As expected, error rates increased amongst more distant genomes (Fig. 4). Unexpectedly, the error rates seem to be similar among all programs, with a slight tendency to be lower for the faster programs than for blastp. The mmseqs results consistently showed the lowest error rate estimates, followed closely by the results obtained with diamond. These results suggest that the quality of orthologs remains as high, if not better, when using software that produces results faster than blastp. The reason why diamond and mmseqs showed the best quality could be that these two programs use fast implementations of the Smith-Waterman algorithm to produce their final alignments.

**Figure 4.**
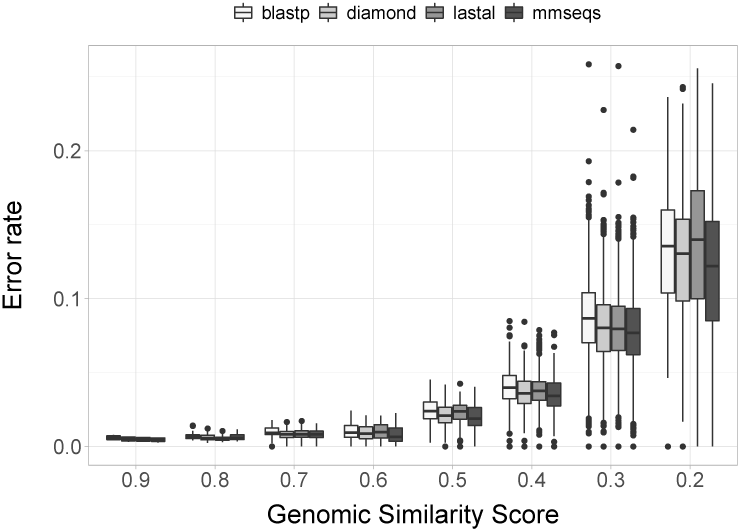
Error rate estimates. All error rate estimates are very close, indicating that using any of the fast programs would not add errors beyond what would be obtained with blastp.

## 4 Concluding remarks

The results above suggest that diamond might be the best alternative to obtain RBH in terms of the speed, sensitivity and quality. As stated, we ran the program using the sensitive option. However, the more-sensitive option seems to run as quickly and to produce very similar results. If much better accuracy is required, mmseqs, with 5.7 sensitivity, produced the most similar results to those obtained with blastp, still in much less time. Our results also showed that the faster programs produced results with very similar error rate estimates as blastp, with diamond and mmseqs showing slightly lower estimates.

Note that our results were obtained comparing prokaryotic genomes. Improvements in speed and sensitivity become more important with larger databases, such as those involving eukaryotic genomes. Thus, the performance of the software might differ when comparing large databases.

## Competing interests

The authors declare that they have no competing interests.

## Author’s contributions

Text for this section …

## Acknowledgements

This work was supported by a Discovery Grant to GM-H from The Natural Sciences and Engineering Research Council of Canada (NSERC). JH-S was supported by a fellowship from Mexico’s Consejo Nacional de Ciencia y Tecnología (CONACYT).

## Notes

https://github.com/Computational-conSequences/SequenceTools

### Competing Interest Statement

The authors have declared no competing interest.

### Summary of Updates

Changed format for a different journal. Added some results using different sensitivity options for mmseqs. Presented all the runtimes together.

